# Light-driven ATP production promotes mRNA biosynthesis inside hybrid multi-compartment artificial protocells

**DOI:** 10.1101/2020.02.05.933846

**Authors:** Emiliano Altamura, Paola Albanese, Roberto Marotta, Francesco Milano, Michele Fiore, Massimo Trotta, Pasquale Stano, Fabio Mavelli

## Abstract

The construction of energetically autonomous artificial protocells is one of the most urgent and challenging requirements in bottom-up synthetic biology. Here we show a hybrid multi-compartment approach to build Artificial Simplified-Autotroph Protocells (ASAPs) in an effective manner. Chromatophores obtained from *Rhodobacter sphaeroides* accomplish the photophosphorylation of ADP to ATP functioning as nanosized photosynthetic organellae when encapsulated inside artificial giant phospholipid vesicles. Under continuous illumination chromatophores produce ATP that in turn sustains the transcription of a DNA gene by T7 RNA polymerase inside ASAPs. Cryo-EM and time-resolved spectroscopy were used for characterizing the chromatophore morphology and the orientation of the photophosphorylation proteins, which allow high ATP production rates (up to ~100 ATP/s per ATP synthase). mRNA biosynthesis inside individual vesicles has been determined by confocal microscopy. The hybrid multi-compartment approach here proposed appears at the same time convenient and effective, and thus very promising for the construction of full-fledged artificial protocells.

---

Bottom-up synthetic biology foresees the construction of artificial protocells as the main ambitious goal^1–6^. Following the pioneer phase^7–10^, current challenges refer to the implementation of more complex behaviour, such as chemical signalling^11–14^, population-level dynamics^15,16^, self-reproduction^17–19^, and autonomous generation of energy^20–22^.

Metabolism is the ensemble of out-of-equilibrium processes within living organisms and includes – but is not limited to – biopolymer synthesis, sensing and motility. It is energy consuming and requires ATP for its fuelling. ATP is generated in vivo by (i) photosynthetic phosphorylation, (ii) oxidative phosphorylation, and (iii) substrate-level phosphorylation. Photosynthetic and oxidative phosphorylations share a common design that requires the establishment of a proton gradient and a difference of chemical potential across the lipid membranes of cells which ultimately triggers the membrane-bound enzymatic complex ATP synthase. In photosynthesis, the gradients are generated by the absorption of light, which needs to be transduced into reduced chemical species by a rather complex dedicated enzymatic apparatus.

Photosynthetic anoxygenic bacteria are ancient photosynthetic organisms that appeared on Earth long before any oxygenic species and possess a simple apparatus for proton translocation. In the particular case of the purple non-sulfur bacterium *Rhodobacter* (*R.*) *sphaeroides* it consists in a minimal set of well-characterized integral membrane enzymes: the reaction center (RC), the ubiquinol:cytochrome c oxidoreductase (bc1) and the ATP synthase (ATP*syn*). The first two complexes are engaged in a photoinduced cyclic electron transport and transmembrane proton translocation involving the coenzyme Q (Q/QH_2_) and cytochrome c_2_ (cyt^2+^/cyt^3+^) redox pools. ATP*syn* eventually converts the proton gradient into ATP molecules, via ADP phosphorylation (Figure 1a).

**Figure 1.**
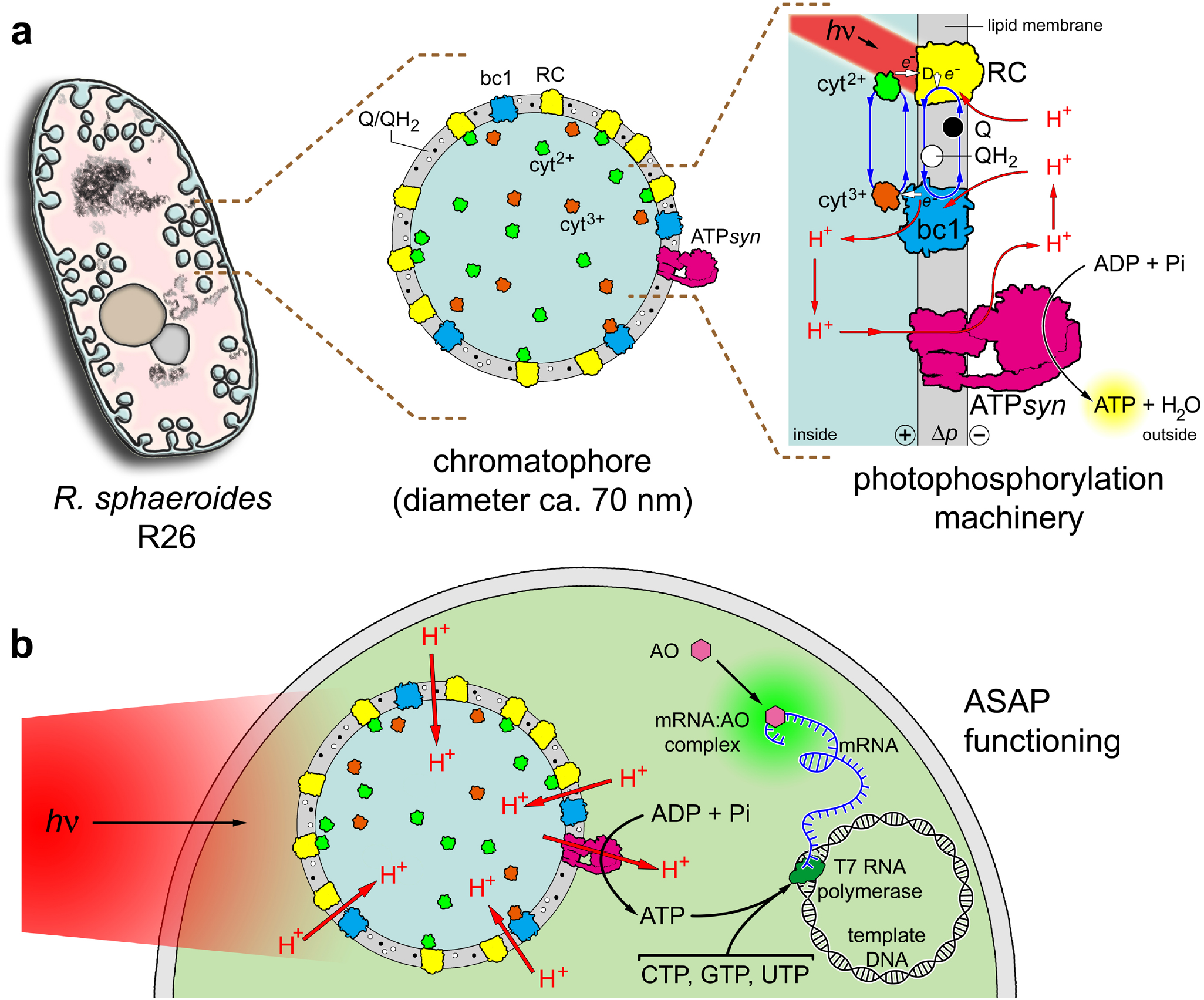
Chromatophore-containing ASAPs. **(a)** *R. sphaeroides* (R26 strain) is a rod-shaped, Gram-negative, purple non-sulfur bacterium that, in absence of oxygen, develops intracytoplasmatic membranes either as invaginations either as organellae-like vesicles, housing the photophosphorylation apparatus^47^. Chromatophores are 60-70 nm closed vesicles derived from the lysis of *R. sphaeroides* cells, typically via French pressing or sonication. Their membrane also contains light-harvesting complex 1 (LH1, not shown for sake of clarity), reaction center (RC, yellow), ubiquinol:cytochrome *c* – oxidoreductase (bc1, blue), ATP synthase (ATP*syn*, purple) in inside-out orientation when compared with the photosynthetic membrane. The chromatophore lumen contains periplasmic solutes, in particular the cytochrome *c*_2_ (here shown in both oxidation states, *i.e.*, reduced cyt^2+^, green; and oxidized cyt^3+^ orange), whereas the chromatophore membrane hosts the ubiquinone pool (oxidized Q, black disks and reduced QH_2_, white disks). The photophosphorylation mechanism starts with a light-induced oxidation of a bacteriochlorophyll dimer (D) in RC. The photogenerated electron travels through the RC and reduces the bound Q that, in two steps, forms QH_2_ (two protons are extracted from the outer solution). The oxidized dimer is reduced by cyt^2+^ present in the chromatophore lumen. The bc1 complex restores the initial conditions by transferring electrons from QH_2_ to cyt^3+^, with a concomitant translocation of protons from the outer solution to the chromatophore lumen. The resulting proton electrochemical gradient (positive inside) is exploited by the ATP*syn* rotatory mechanism to catalyse the thermodynamically uphill conversion of ADP + Pi into ATP (ca. 100 ATP/s per ATP*syn*), produced in the outer solution. Chromatophores act as ATP-producing organellae when illuminated by 860 nm light. **(b)** ASAP functioning. Chromatophores are encapsulated inside ~20 μm giant phospholipid vesicles made of POPC (1-palmitoyl-2-oleoyl-*sn*-glycero-3-phosphocholine) and illuminated to generate a proton motive force (ca. 130 mV) across the membrane. Co-encapsulated ADP, Pi, GTP, CTP, UTP, T7 RNA polymerase (dark green) and a DNA template give rise to an out-of-equilibrium system of coupled reactions, namely, photophosphorylation and DNA transcription. The biosynthesized mRNA is revealed by acridine orange (OA, pink), which binds it forming a green-fluorescent complex.

In the perspective of exploiting the *R. sphaeroides* apparatus for producing ATP inside artificial protocells upon light irradiation, we have recently started a systematic investigation about these sorts of systems that we call photoactive “Artificial Simplified-Autotroph Protocells” (ASAPs) (see Text S1 for the used terminology). We have previously developed a single-compartment approach, consisting of vesicles functionalized with >90% oriented protein complexes able to transduce light energy in chemical energy in form of a proton gradient^23, 24^. Here we report a novel Hybrid Multi-Compartment Approach (HyMCA), consisting in the entrapment of nanosized biological vesicles (< 100 nm) inside a giant artificial vesicle (> 1 μm). In particular, we have employed *R. sphaeroides* “chromatophores” (closed cytoplasmic membrane fragments, see below) as light-driven ATP-synthesizing organellae trapped inside phospholipid giant unilamellar vesicles (GUVs). The nascent ATP has been channelled into DNA transcription, here catalysed by T7 RNA polymerase in the presence of the other three nucleotide triphosphates (CTP, UTP and GTP) to form mRNA strands ultimately revealed by the fluorescent marker acridine orange (Figure 1b).

To this aim we have firstly obtained functional chromatophores from *R. sphaeroides* and characterized them with respect to their morphology, composition, and light induced ATP synthase activity. Next, we have encapsulated the chromatophores and the other required components inside GUVs by the highly efficient droplet transfer method, demonstrating the successful intravesicle production of ATP. Finally, we have coupled ATP synthesis with DNA transcription in order to produce a messenger RNA. In this way we assembled “ASAPs” (see Text S1) to perform ATP-sustained metabolic transformations and therefore mimicking key biological reactions in a realistic way.

From the technological viewpoint, our hybrid approach provides a convenient and efficient route to the construction of artificial protocells of higher complexity, for example when ATP-driven processes and networks are required. This is made possible thanks to the built-in energy transduction capacity of chromatophores.

## Obtaining functional chromatophores from *R. sphaeroides*

The photosynthetic phosphorylation apparatus of *R. sphaeroides* R26 resides in the cytoplasmic membrane as a set of pigments, integral membrane proteins and protein complexes (light harvesting complexes LH1, RC, bc1, ATP*syn*). Cytochrome *c*_2_ in the periplasm and membrane quinones are also necessary to the photoactivity of the protein apparatus, being both involved as electron carriers. The cytoplasmic membrane is actually extended into the cell as hundreds of vesicle-like invaginations, with the F_1_ subunits of ATP synthase facing the cytoplasm. When the cell is broken, these invaginations pinch off, yielding sealed vesicles the chromatophores. These membrane vesicles generally have the photophosphorylation machinery in their membrane with an inside-out orientation and encapsulate the cytochrome *c*_2_ in their internal milieu (Figure 1a). Accordingly, purified chromatophores are able to produce ATP (in their outer solution) when irradiated with Near InfraRed (NIR) light in the presence of externally added ADP and inorganic phosphate (Pi)^25,26^. To obtain high yield of chromatophores with the desired orientation of photophosphorylation machinery, we have employed an extraction procedure based on direct cell lysis (e.g., by a single French press passage, Text S2), followed by centrifugation steps^27,28^ (Figure 2a).

**Figure 2.**
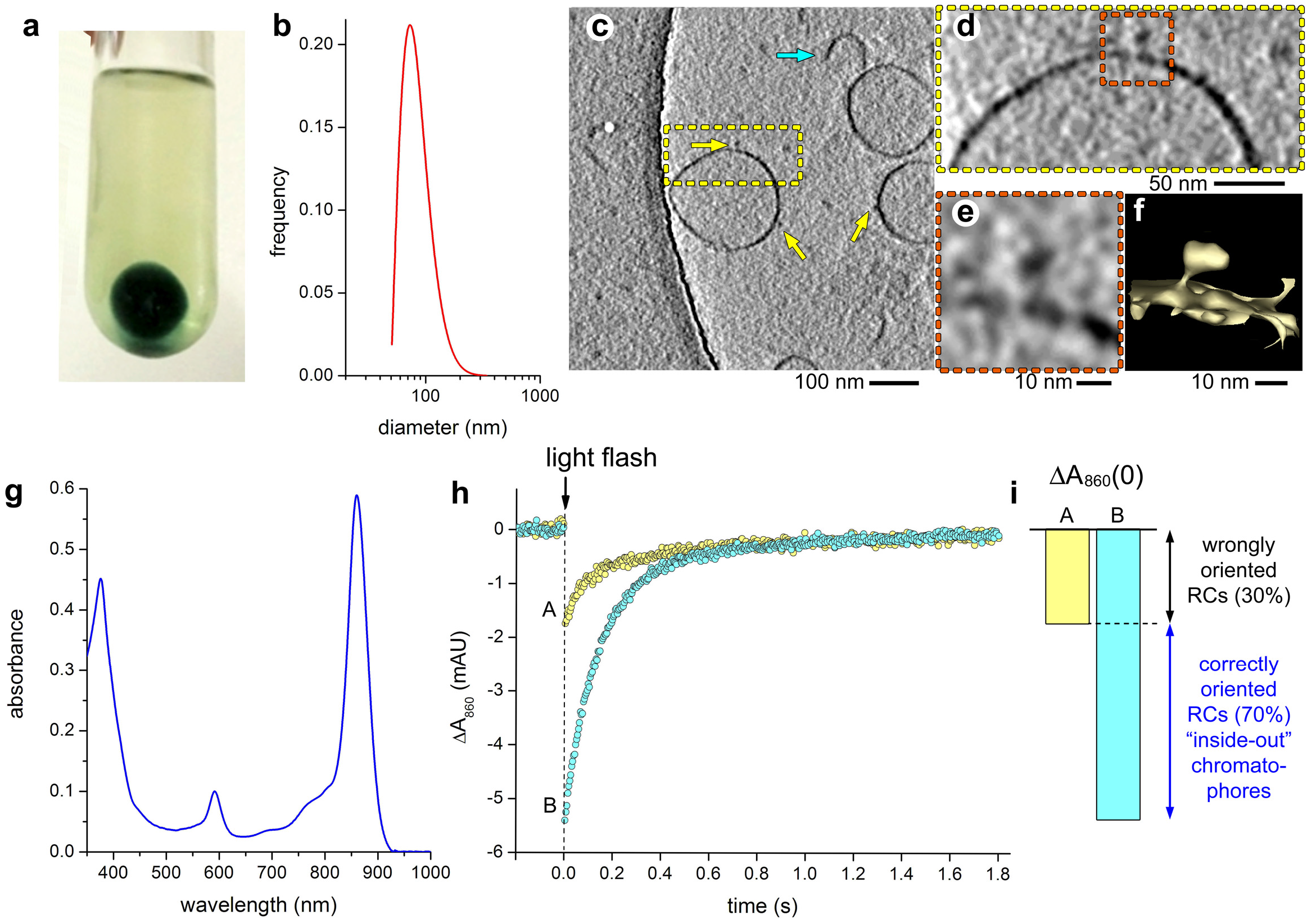
*R. sphaeroides* chromatophores. **(a)** Green chromatophores produced by French pressing *R. sphaeroides* cells and pelleted by ultracentrifugation. **(b)** Number-weighted size distribution of chromatophores as obtained by DLS (the curve refers to an average of three measurements of an individual sample; mode: 68 nm; mean ± SD: 80 ± 23 nm). **(c,d)** Cryo-EM tomographic slice of chromatophores; yellow arrows indicate the clearly visible ATP synthase in their equatorial plane, cyan arrow indicates an open membrane fragment; additional images in Figure S1. **(e)** Magnification of a single ATP synthase (13 nm wide, 21 nm long); additional images in Figure S2. **(f)** 3D-Tomographic reconstruction of ATP synthase shown in **(e)**. **(g)** UV-Vis absorption spectrum of purified chromatophores; note the peak at 860 nm, due to bacteriochlorophylls, that serves as a measure of chromatophore concentration. **(h)** Kinetics of charge recombination taking place within the RC. The amplitude of the 860 nm peak suddenly decreases upon actinic light irradiation (saturating flash, indicated by the arrow), due to charge separation. Next, a slow increase of the 860 nm absorbance is observed (completed in about two second), due to charge recombination. The two curves refer to an individual chromatophore sample before (yellow points, A), and after treatment with 1% w/v LDAO (cyan points, B). **(i)** Comparison between Δ*A*_860_(0) (bar A) and Δ*A*_860_(0),_LDAO_ (bar B) and their meaning in terms of RC orientation in the chromatophores. The bars represent the values reported in Figure 2h and refer to an individual chromatophore sample. Size bars represent 100 nm **(c)**, 50 nm **(d)**, 10 nm **(e,f)**.

The chromatophore diameter was measured by dynamic light scattering (mode 68 nm; mean 80 ± 23 nm, Figure 2b), while their zeta potential is −30.9 ± 0.6 mV (mean ± SD), as expected for the presence of ~ 60 mol% of anionic phospholipids in the *R. sphaeroides* membrane^29, 30^.

Cryo-EM and cryo-electron tomography confirm the presence of closed spheroidal chromatophores although some elongated vesicles and open membrane fragments have been also observed (Figure 2c). Direct inspection of cryo-EM images clearly reveals the presence of F_0_F_1_ ATP synthase complex in the functional orientation (the F_1_ subunit pointing outward) on the membrane of closed chromatophores (Figure 2d-e and Figure S1). On average, we found 1.6 ATP*syn* per chromatophore, all being outward-oriented and thus able to produce ATP in the external solution. In no case inward-oriented ATP synthases were found. Image analysis gives a surface density of ca. 50 ATP*syn*/μm^2^. The tomographic 3D reconstruction (Figure 2f) renders the characteristic ATP synthase structure, expectedly, as a ball-on-a-stick complex 13.2 ± 1.2 nm wide and 21.2 ± 1.9 nm long (Figure S2).

In order to be functional (photo-active), chromatophores with inside-out morphology (Figure 1a) should also contain enough amount of cyt^2+^ in their lumen. We have determined the ratio between functional chromatophores and all other types of non-functional particles and fragments that arise from the French press treatment by a flash light excitation experiment^23^ (details in Text S2-S3). In this assay the RC spectral change at 860 nm (ΔA_860_), due to a photo-induced charge-separated state, is recorded immediately after the flash (see Figure 2gh). Because cyt^2+^ reduces RC in the charge-separated state and quenches ΔA_860_, the measured ΔA_860_(0) is proportional to the concentration of RCs not instantaneously reduced by cyt^2+^, *i.e*., RCs in non-functional particles. When the detergent LDAO is added, functional chromatophores and all other particles are converted into micelles. Functional chromatophores release the entrapped cyt^2+^ in the medium, where it is highly diluted (10,000-fold). In these conditions, all RCs contribute to ΔA_860_(0),_LDAO_ because no RC can be reduced efficiently by cyt^2+^. From the comparison between ΔA_860_(0) and ΔA_860_(0),_LDAO_ (Figure 2i) it can be determined the concentration of 6.3 μM of RCs in functional chromatophores (Figure 1a), representing about 70% of the total RC content. Moreover, the concentration of functional chromatophores (~10^15^/L when OD = 1) can be also calculated thanks to the known average RC surface density in the *R. sphaeroides* chromatophores (ca. 2,300 RC/μm^2^)^31, 32^.

In conclusion, by combining spectral analysis, DLS and cryo-EM images, we defined the average structure and composition of the chromatophores used in this study (Table S1). Importantly, we have shown that ready-to-use “functional” chromatophores can be obtained by a single French press passage with high yield (≥ 70%, see Text S2). As it will be shown below, they are capable of transducing light energy in biochemical energy with a very high ATP synthase activity (ca. 20 μmol_ATP_ min^−1^ mg^−1^_ATP*syn*_).

## Assaying ATP production by chromatophores in bulk

We assayed chromatophores in bulk for their capacity of producing ATP on their external medium upon continuous illumination, in the presence of externally added ADP and Pi. The production of ATP would confirm that chromatophores are fully functional and operate according to the scheme indicated in Figure 1a. The amount of synthesized ATP has been quantified by the luciferin-luciferase bioluminescence assay, according to a calibration line. Preliminary experiments, summarized in Text S4 and Figure S3, lead us fixing optimal conditions for the assay, *i.e.*, ~6 × 10^16^ chromatophores/L (OD = 50), 2 mM ADP, 100 mM Pi (pH 8.0), and 3 minutes NIR light irradiation (860 nm). Not surprisingly, in absence of a process that consumes ATP, the ADP-to-ATP conversion is high but not complete (60%). The calculated reaction quotient [ATP]/([ADP][Pi]) ≅ 15 M^−1^ is compatible with a proton motive force of ca. 130 mV (positive inside chromatophores) if we consider 3 translocated protons per ATP molecule^34^. When 2 mM ADP is employed, the ATP production rate almost reaches the maximum (99.5% of *V*_max_, considering *K*_M,ADP_ = 10 μM for the ATP synthase^35, 36^). The measured production of 1.2 mM ATP in 3 minutes corresponds to an ATP synthase turnover number of ~100 s^−1^ (or higher), in good agreement with the highest reported values (50 s^−1^ ^37^; or 90 s^−1^ ^38^) (see Text S5 for a commentary). When compared to previous artificial protocell reports^20,21^, which are based on other types of ATP synthase-bearing artificial organellae, chromatophores perform 12-23× better.

The components of the photophosphorylation machinery embedded in the chromatophore membrane are highly functional because they reside in their own physiologic lipidic microenvironment. Moreover, the results demonstrate that chromatophores are endowed with sufficient and well-oriented bc1 complex, as well as with enough amount of Q/QH_2_ pool in the membrane and a sufficient amount of cyt^2+^ in the lumen.

## Assembling ATP-producing multi-compartment protocells

The construction of chromatophore-containing protocells is conveniently pursued using GUVs prepared by the droplet transfer method using POPC as phospholipid^39, 40^ (Figure 3a). A suspension of chromatophores is firstly emulsified in a phospholipid-containing mineral oil solution and the resulting water-in-oil droplets – which are stabilized by a phospholipid monolayer – are then transferred to an underlying aqueous phase by centrifugation. When the water-in-oil droplets cross the oil/aqueous interface, see Figure S4, they become covered by a second phospholipid monolayer, and generate chromatophores-containing protocells. The chromatophore encapsulation inside protocells is straightforward, leading to stable protocells whose typical appearance is shown in Figure 3b. Note that direct chromatophore imaging by optical microscopy is prevented by their small size, however their presence is revealed by lipophilic Nile Red staining.

**Figure 3.**
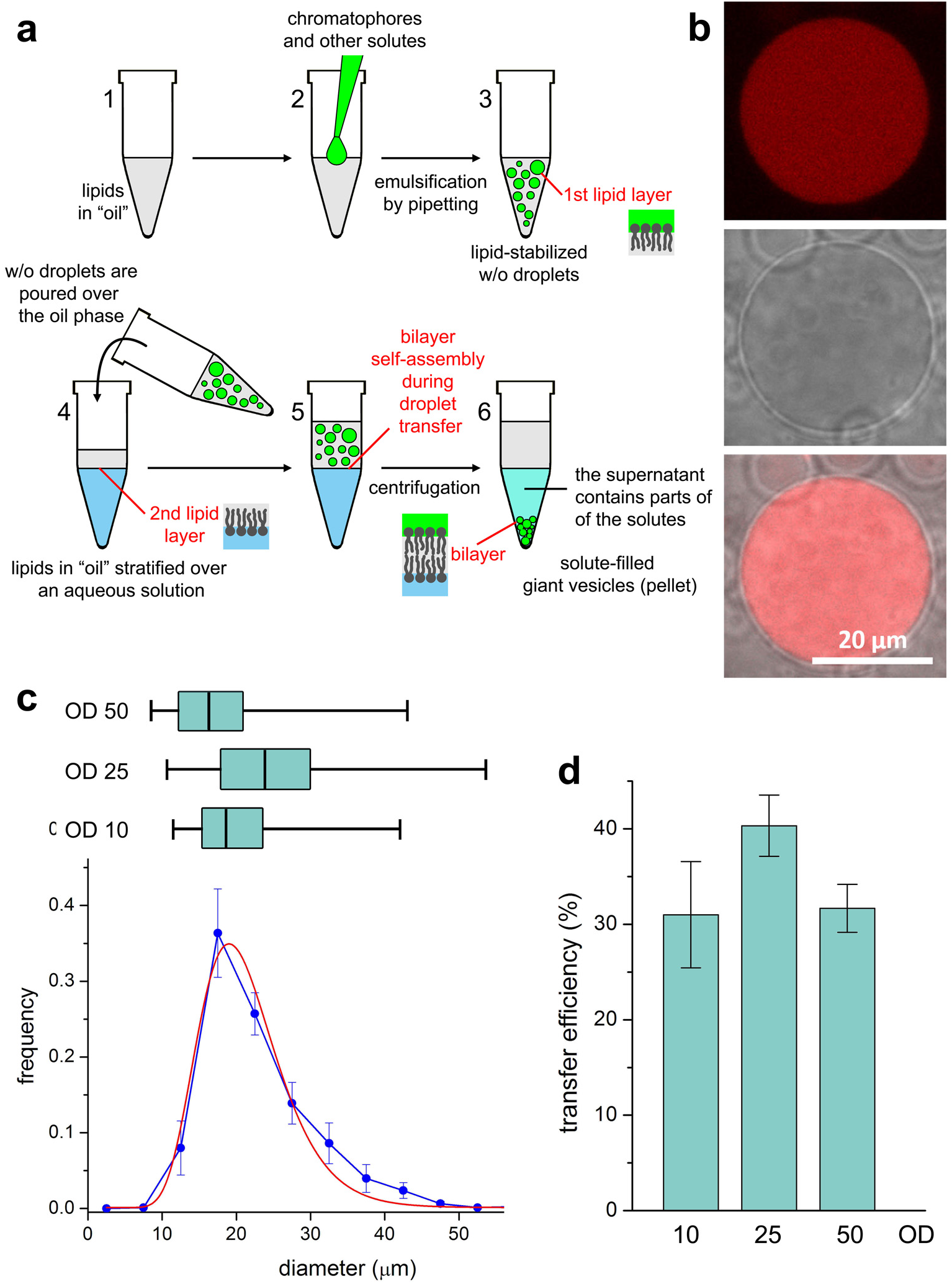
Preparation of multicompartment protocells. **(a)** The droplet transfer method consists in two steps. In the first, lipid-stabilized w/o droplets are prepared by mechanical emulsification of a chromatophore-containing aqueous solution in a lipid-containing mineral oil solution. Chromatophores and all water-soluble solutes are entrapped inside the w/o droplets. Next, the w/o droplets are gently poured over another vial consisting of a lipid-in-oil solution stratified over an aqueous phase. GUVs are formed when the w/o droplets cross the second lipid monolayer at the oil-water interface. Chromatophores and water-soluble compounds are found inside GUVs. The droplet transfer is not 100% efficient; some w/o droplets break during the centrifugation, releasing their content in the bottom aqueous solution. **(b)** Appearance of multicompartment protocells (fluorescent channel, top; bright field, center; overlay, bottom). Chromatophores have been stained by the hydrophobic fluorescent dye Nile Red. The size bar represents 20 μm. **(c)** Protocell size distribution (blue points) obtained by image analysis of protocells (*n* = 948, mean ± SE). The red curve represents the best-fit log-normal distribution (mean: 20.5 μm). The horizontal boxes represent the three protocell size distributions obtained by varying the concentration of entrapped chromatophores (OD 10, *n* = 234; OD 25, *n* = 171; OD 50, *n* = 194); the two extreme marks indicate the limits of the distribution, the box marks indicate the first, the second, and the third quartiles. The three medians are not significantly different according to the Kruskal-Wallis test for non-normal distributions. **(d)** Comparison between the transfer efficiency as function of the concentration of entrapped chromatophores (OD 10, 25, 50). Bars represent the transfer efficiency (mean ± SD, *n* = 3 independently prepared samples) as obtained by measuring the concentration of chromatophores released in the aqueous solution just after the droplet transfer. The three means are not significantly different according to the one-way ANOVA test.

The protocell size distribution and formation efficiency (i.e., fraction of water-in-oil droplets successfully transformed into GUVs) resulted not very sensitive to the chromatophore concentration in their lumen, confirming the robustness of the droplet transfer method (Figure 3c-d). Our procedure leads to about 2 × 10^10^ protocells /L (diameter ~20 μm), with ~35% efficiency. Each 20 μm protocell contains ~48,000 chromatophores (OD 10) accounting for less than 1% of the protocell inner volume. The remaining part, the protocells ‘free volume’, is the locus of ATP production and its further utilization by other reactions (Figure 1b). The average features of a 20 μm protocell are summarized in Table S2.

Firstly, we asses the capacity of light induced proton translocation of encapsulated chromatophores. By co-encapsulating pyranine, a membrane impermeable pH-sensitive dye, we measured an increase of pyranine fluorescence upon continuous NIR irradiation (Figure S5). This increase is ascribed to the rise of pH in the protocell free volume. This is expected when protocells are prepared without ADP and Pi (Figure 1b), because illuminated chromatophores just translocate protons from the protocell free lumen volume to their interior, developing a pH gradient and an electrochemical membrane potential.

Protocells prepared with all required components were then assayed for ATP production. In the presence of ADP and Pi the photo-redox reactions can cycle very efficiently, because the pH gradient and the electrochemical potential are continuously generated by light absorption and thereafter dissipated by ATP synthase. The quantification of intra-protocell ATP (overall concentration, Text S6) can be carried out by the luciferin-luciferase bioluminescence assay after releasing ATP from the protocell interior. Preliminary experiments discouraged detergent-based lysis (Figure S3c) and also indicated minor interference of the sugars present inside and outside the protocells. Therefore, by freezing-and-thawing the sample, ATP has been released from protocells and quantified (Figure S6).

The measured average concentration of intra-protocells ATP (~38 μM) definitely demonstrates that chromatophores act as ATP-producing photosynthetic organellae inside a larger compartment, as required by our design. Note that this experiment does not determine the maximal ADP-to-ATP conversion because ATP synthase, functioning in reverse, partially hydrolyses ATP during the freeze-and-thaw steps.

## Photosustaining mRNA biosynthesis in ASAPs

Having shown that protocells autonomously produce ATP from photon energy, we finally added an anabolic module to the system, in particular the DNA transcription, calling the resulting structures ASAPs (see Text S1). ASAPs were built by including in the protocell free lumen volume ADP, Pi, a linearized plasmid as DNA template (pTRI-Xef carrying the 1.85 kb *xenopus elongation factor 1α* gene under T7 promoter), the enzyme T7 RNA polymerase, and GTP, UTP, CTP. Figure 1b summarizes the whole process. The entire reaction network functions out-of-equilibrium thanks to vectorial chemistry across the chromatophore membrane, entirely driven by the primary photophysical process of charge separation within RC.

The fluorescent dye acridine orange (AO) was added to ASAPs in order to evidence the 1.89 kb mRNA produced from the coupled photophosphorylation-transcription reactions. AO binds both DNA and mRNA, emitting light in the green region (peak at 530 nm, see Figure S7). Because the DNA concentration is constant, the variation of AO fluorescence is only due to the progress of transcription. A series of confocal fluorescence micrographs are shown in Figure 4a and the fluorescence versus time profile is reported in Figure 4b, obtained by quantitative image analysis of ASAP micrographs per each time interval. The accumulation of mRNA inside ASAPs is clearly detected by the sigmoidal increase of the AO:mRNA complex fluorescence. The photophosphorylation reaction is driven out-of-equilibrium both for the continuous actinic illumination and for the transcription reaction. Consequently, ADP is converted completely to ATP, and the latter is incorporated in the nascent mRNA chain. Because ATP is the limiting reagent for T7 RNA polymerase, the maximal amount of mRNA will be ~350 nM, because the adenine base counts around 30% in this specific mRNA.

**Figure 4.**
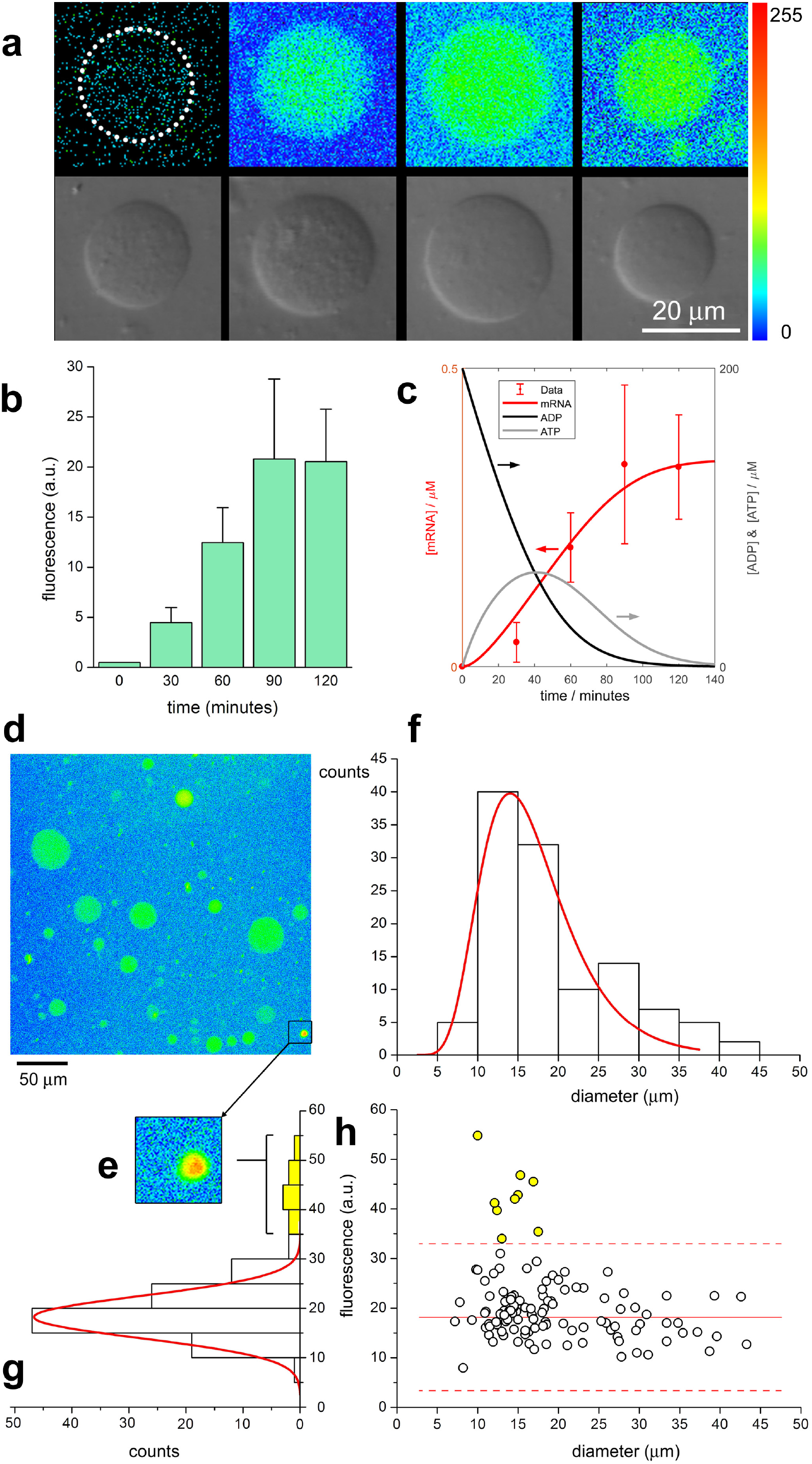
mRNA biosynthesis inside ASAPs. **(a)** Typical appearance of mRNA-biosynthesizing ASAPs after 30, 60, 90 and 120 minutes of illumination and incubation at 37 °C (fluorescent channel, top; bright field, bottom). Fluorescence is due to the complex between acridine orange and mRNA. Images are given in a coded-colour scale, shown on the right (different colours represent different fluorescence intensities). The size bar represents 20 μm. **(b)** Increase of fluorescence inside ASAPs as determined by image analysis over a single population of ASAPs (30 minutes, *n* = 43; 60 minutes, *n* = 121; 90 minutes, *n* = 113; 120 minutes, *n* = 72); bars represent the mean ± SD. **(c)** Calculated time-dependent profiles of ADP (black), ATP (grey), and polymerised ATP (in form of mRNA, red) according to a minimal two-reaction model described in the Text S7. **(d)** Diversity among mRNA-biosynthesizing ASAPs made evident by a wide-field image (coded-colour scale). **(e)** Magnification of a highly fluorescent ASAP. **(f)** ASAP size distribution of data points shown in (h); the red curve represents the best-fit log-normal distribution. **(g)** Fluorescence distribution of data points shown in (h); the red curve represents the best-fit normal distribution matching the major distribution peak; bars outside the normal distribution are marked yellow. **(h)** Dot-plot representing the size and the internal fluorescence of ASAPs after 90 minutes illumination. ASAPs with high fluorescence are marked yellow. The continuous red line and the two dashed red lines represent the average and the boundaries of a sub-population of ASAPs (with a grey background) whose fluorescence is normally distributed.

Intrigued by the sigmoidal profile shown in Figure 4b we designed a minimal kinetic model to explain such behaviour. The model is based on two coupled enzymatic reactions: (1) ADP + Pi → ATP, catalysed by ATP synthase, and (2) ATP + other nucleotides → mRNA + PPi, catalysed by T7 RNA polymerase in the presence of the DNA template (details in Text S7). The numerical integration of the corresponding Michaelis-Menten-type differential equations^41,42^ produces the kinetic profiles of ADP, ATP and ‘polymerised ATP’ (*i.e.*, adenine bases in the mRNA chains), which fit quite well the experimental data (Figure 4c). The best-fit values of the turnover numbers and the Michaelis constants result to be 13 s^−1^ and 110 μM for ATP synthase, and 0.12 s^−1^ and 52 μM for T7 RNA polymerase, in satisfactory agreement with reported values (Table S3).

ASAPs display non-negligible variability, as evident from wide-field imaging (Figure 4d). The sample contains exceptionally fluorescent ASAPs (Figure 4e). The variability arising from intra-ASAPs reactions appears evident in Figure 4f-h, where a fluorescent-*vs*-size dot plot disentangles the two variables, revealing the existence of a sub-population of ASAPs displaying especially efficient internal reactions. In particular, some ASAPs have high fluorescent values (i.e., high mRNA concentration) that fall outside the Gaussian peak (Figure 4g) matching the peak of the fluorescence distribution. Highly fluorescent ASAPs are typically small. These variations can be due to an anomalous stochastic partition of the solutes and of chromatophores encapsulated in the very moment of ASAP formation, as reported in other artificial protocell studies prepared (reviewed in^43^). Such an intriguing phenomenon would generate diversity among ASAPs and would produce, ultimately, large variations of the reaction kinetics, product concentration, and fluorescence. To test the hypothesis of anomalous solute partition, a useful approach is based on the comparison between experiments and computer simulations, where populations of compartments are generated in silico according to various solute partition laws^42^. This sort of study is outside the scope of the present work and it will be investigated in a follow-up report.

## Conclusions

We have presented a prototype of a photoactive Artificial Simplified-Autotroph Protocell (ASAP), made of GUVs that encapsulate *R. sphaeroides* chromatophores able to transduce light energy in chemical energy. Chromatophores act as natural photosynthetic organellae enclosed in an artificial protocell converting ADP and Pi into ATP at the sole expense of light energy. This sort of energy transduction can play a central role for the engineering of long-lasting artificial protocells^44^ capable of generating the required high-energy compounds and thus fuelling internal processes. Here we have focused on DNA transcription, but other processes can be envisaged.

Two considerations are specific about this work. The first is the hybrid design. As an alternative to a total bottom-up approach, a HyMCA takes advantage of ready-to-use components borrowed from biological organisms (natural or engineered ones). Chromatophores are *par excellence* the simplest ATP-producing building blocks and can be employed in modular design of artificial protocells. Our hybrid design can be further expanded to include other easy-to-extract natural organellae (chloroplasts, mitochondria), increasing the versatility of bottom-up approaches and therefore accelerating the possible implementation of artificial protocells in real-world applications. A few works have already reported similar combinations^45,46^. The second consideration refers to the out-of-equilibrium and energy-transducing features of living systems, here mimicked in a genuine way. The photoactive ASAPs presented in this work produce the key intermediate ATP by exploiting an environmental source of energy (NIR light) and immediately consume it for sustaining the polymerization of mRNA. The ATP-producing and ATP-draining reactions both concur to the establishment of the out-of-equilibrium dynamics, highlighting the essence of cellular metabolism: the energy flow through intermediate molecules.

Finally, we remark that the primary energy source in our design – *i.e.*, light – offers specific advantage due to its easy quantitative/qualitative manipulation, and points to interesting future opportunities in artificial protocell control.

## Acknowledgements

This work was partially supported by CSGI (Consorzio Sistemi a Grande Interfase). The authors thank Dr. Lucia D’Accolti of University of Bari for financial support, Prof. Gerardo Palazzo and Dr. Luigi Viggiano of University of Bari for DLS analysis and for the MegaScript™ T7 transcription kit, respectively. Dr. Massimo Lasorsa and Dr. Nicola Altamura of National Research Council (CNR) of Bari are acknowledged for supplying of several chemicals.

## Online Methods

### Production of *R. sphaeroides* chromatophores

Cells of *R. sphaeroides* (strain R26) were purchased from the Deutsche Sammlung von Mikroorganismen und Zellkulturen GmbH (Braunschweig, Germany) (DSMZ, #2340). Bacterial cells were grown in 1 L of DSMZ medium (Tables S4-S6), pH 6.9 at 25 °C, first oxygenically in the dark for 12 hours, then photoheterotrophically under anaerobic conditions exposing the cell culture to continuous light with a 60 W tungsten filament light bulb placed at 25 cm from the center of the bottles. In these conditions the anoxygenic, photosynthetic growth initiates, entering in the exponential growth phase. After three days R26 cultures assume a characteristic intense green/blue colour because of the absence of carotenoids molecules. Harvested cells were centrifuged at 15,300 *g* for 15 minutes at 4 °C, and the pellet was washed twice with 200 mL of phosphate-EDTA buffer (PE-buffer; 5 mM K-phosphate pH 8.0, 1 mM EDTA), and finally resuspended and homogenized using a brush and a potter in 50 mL of PE-buffer. Few flakes of DNase (Sigma, #DN25) and 5 mM MgSO_4_ were added. After 15 minutes incubation at 4 °C, the cells were disrupted by a single French press passage operating at 16,000 psi (1,100 bar) and 4 °C. The sample was centrifuged at 15,300 *g* for 15 minutes at 4 °C, to remove the cell debris and heavy fragments (in the pellet), while the supernatant (chromatophores) was further ultracentrifuged at 140,000 *g*, for 120 minutes at 4 °C. The pelleted chromatophores were resuspended in PE-buffer (~5 mL), then centrifuged again at 15,300 *g*, 4 °C, for 15 minutes to remove residual debris and heavy fragments. The supernatant contains chromatophores whose optical density at 860 nm (OD) is typically ~50. If needed, chromatophores can be concentrated by pelleting and resuspension.

### Cryo-EM imaging and tomographic 3D reconstruction

Frozen hydrated samples were prepared by applying a 3 μL aliquot to a previously glow discharged 200-mesh Quantifoil 1/2 holey carbon grid (Ted Pella, USA). Before plunging the grid into liquid ethane, the grid was blotted in a chamber at 4 °C and 100% humidity using a FEI Vitrobot Mark IV (FEI company, the Netherlands). The particles were imaged using a Tecnai F20 microscope (Thermo Fisher Scientific - US), equipped with a Schottky Field Emission electron source operated at an acceleration voltage of 200 kV, a US1000 2k × 2k Gatan CCD camera and a FEI retractable cryo box to limit water sublimation from the vitrified sample and its recrystallization on the specimen surface. For the cryo-electron tomography the tilt series were collected by tilting the vitrified sample over ± 60° with the following tilt sequence: starting from 0° to ± 48 with a tilt step of 3°; then, from ± 48° to ± 60°with a tilt step of 2°. The cryo-EM imaging was carried out at a final object pixel of 3.6 Å, with a total dose of ∼60 e^−^/Å^2^ in order to limit specimen damage. Computation of the tomogram and isosurface based segmentation was carried out with the IMOD software package using a WBP-based algorithm. [M1].

### Dynamic light scattering (DLS) and zeta-potential measurements

DLS measurements were performed using a Zetasizer-Nano S from Malvern operating with a 4 mW He-Ne laser (633 nm) and a fixed detector angle of 173° (non-invasive backscattering geometry) and with the cell holder maintained at 25 °C by means of a Peltier element. Data were collected leaving the instrument free to optimize the instrumental parameters (attenuator, optics position and number of runs) according to house-built algorithm, leading to the determination of the *z*-average size, the size distribution, and the zeta-potential of chromatophores. Note that the obtained intensity-weighted size distribution function, that overestimates the larger particles, was finally converted into a number-weighted size distribution. For all measurements (in triplicates), two chromatophores concentrations were measured (OD 0.6 and 0.06), without major differences between the results.

### RC quantification, orientation, and sealing assay

RC content and orientation can be assayed by flash photolysis experiment [M2]. Flash photolysis consists in promoting the reagents in an excited state by using an intense light impulse and then in monitoring the spectral changes thanks to a lower-intensity secondary light, which is not interfering with the system. In the case of RC, the light flash induces a charge-separated state (CSS) in all RC molecules, and the spectrophotometer records the time evolution of absorbance at 860 nm, after the flash. Chromatophores (OD 0.6) suspended in 10 mM K-phosphate buffer (pH 7.4) were placed in a 1-cm squared quartz fluorescence cuvette and irradiated by xenon lamp flashes (~100 μs) placed orthogonal with respect to the measuring beam. The absorbance drop at 860 nm (Δ*A*_860_), which mirrors the CSS, was followed in time (for about 2 s). Data were collected onto a digital oscilloscope (Tektronics TDS-3200). A second experiment has been carried out on chromatophores treated with 1% w/v LDAO, which converts chromatophores into in micelles. The difference between the initial Δ*A*_860_ signal in presence and in absence of LDAO is proportional to the amount of RC molecules correctly oriented (*i.e.* with the chromatophore binding site facing the chromatophore lumen). Further details are given in Text S2.

### Quantification of light-induced ATP production by chromatophores in bulk

The luminescence assay solution (10 mL) was freshly prepared before each experiment by mixing MilliQ water (8.9 mL), 20× assay buffer (500 mM tricine pH 7.8, 100 mM MgSO_4_, 2 mM EDTA, 2 mM NaN_3_) (0.5 mL), 100 mM DTT (0.1 mL), 10 mM d-luciferin (0.5 mL), and 5 mg/mL firefly luciferase (2 μL).

Samples were prepared on a 10 μL scale. Chromatophores (OD 50) were mixed with 2 mM ADP and 100 mM K-phosphate buffer (pH 8) and illuminated for 3 minutes in a quartz ultramicrocuvette (Hellma, #105.250) by a 860 nm/2.6 W LED lamp (OSRAM, #SFH 4783), which provide an irradiance of 25 mW/cm^2^, equivalent to about 1.1 × 10^17^ photons/s/cm^2^. ATP quantitation was carried out in the same microcuvette, by adding 90 μL of the assay solution (see above) and measuring the initial value of the developed bioluminescence (560 nm) by means of a Cary Varian Eclipse spectrofluorometer set as luminometer in kinetic mode. Negative control samples, additional assays, and calibration samples (1-10 μM ATP in absence and in presence of chromatophores) have been prepared and measured similarly.

### Preparation of chromatophore-containing protocells by the droplet transfer method

The droplet transfer method was applied [M3], by following an optimised procedure [M2, M4]. In a 1.5 mL Eppendorf tube (tube A), 300 μL of organic phase consisting in 0.5 mM POPC (Lipoid GmbH, Steinhausen, Switzerland) dispersion in mineral oil (Sigma-Aldrich, #M5904), were gently overlaid over 500 μL of aqueous O-solution (200 mM glucose and the same buffer as in the I-solution), in order to create a lipid-covered interfacial region. In another 1.5 mL Eppendorf tube (tube B), 20 μL of I-solution (200 mM sucrose, chromatophores in PE-buffer A, and other solutes when required) were added to 600 μL of 0.5 mM POPC in mineral oil. A water-in-oil (w/o) emulsion was obtained by pipetting repeatedly up and down the mixture for 30 seconds. Next, the w/o emulsion (tube B) was gently poured on top of the organic phase in tube A. Tube A was centrifuged at 2,500 *g* for 10 minutes at room temperature. After the centrifugation, the mineral oil appeared clear and it was removed. The resulting protocells, forming a pellet on the bottom of the tube, were gently collected by direct aspiration with a polypropylene micropipette tip (50 μL). To obtain an oil-free protocells suspension, care was taken to wipe off, with a tissue paper, the small amount of oil possibly collected on the outside of the pipette tip. Protocells were washed twice by centrifugation, supernatant removal, and re-suspension in fresh O-solution to remove non-entrapped substances. When dispersed in a final volume of 50 μL, a typical concentration of 2.2 × 10^7^ protocells/mL was obtained.

Protocells size distribution was measured by confocal microscopy image analysis. The transfer efficiency is defined as the fraction of w/o droplets that are successfully transformed in protocells, thanks to a successful transfer from the oil phase to the O-solution. It can be estimated, under simplifying assumptions, by measuring the entrapment efficiency, *i.e.* the ratio between the amount of chromatophores encapsulated inside protocells and the total amount of chromatophores used for the preparation. Entrapment efficiency is measured by measuring the 860 nm absorbance readings of: (*i*) the O-solution supernatant (*A*_s_) just after the droplet transfer (the supernatant contains non-entrapped chromatophores, whereas protocells are in the pellet); and (*ii*) the same volume of O-solution spiked with 20 μL of chromatophore-containing I-solution (*A*_tot_). The entrapment efficiency (%), and thus the transfer efficiency (%), is calculated as 100 × (1 − *A*_s_/*A*_tot_).

### Evidencing the light-driven generation of transmembrane pH gradient

Protocells containing chromatophores (OD 50) and pyranine (5 μM) were prepared as described, in absence of ADP and Pi. Samples were illuminated directly on the glass slide with a 860 nm/2.6 W LED lamp, for 5 minutes. After every minute of continuous illumination an image of protocells has been recorded (during the acquisition time, the LED lamp was switched off). The negative control sample consisted in protocells filled with pyranine only.

### Quantification of light-induced ATP production by chromatophores inside protocells

Six samples of protocells filled with chromatophores (OD 10), 200 μM ADP and 10 mM Pi were prepared as previously described and pooled together in 90 μL final volume. One at a time, six 10 μL aliquots were illuminated for 10 minutes with a 860 nm/2.6 W LED lamp, then the samples were frozen in liquid nitrogen and thawed in a water bath at 40 °C for 10 times, in order to release ATP from the protocells lumen. Known amounts of ATP (0, 10 and 20 μM) were then spiked into the samples (*i.e.* the standard additions method), followed by the ATP bioluminescence assay as described above.

### Coupled light-induced ATP production and mRNA biosynthesis inside ASAPs

ASAPs containing chromatophores (OD 10), 200 μM ADP, 10 mM Pi and a commercial transcription kit (MEGAscript® T7 Transcription Kit, Ambion™, #AM1334) were prepared as described. The transcription kit contained CTP, UTP, GTP (7.5 mM each), 0.5 μg of pTRI-XEF linear DNA template (which codes for the 1.85 kb *Xenopus elongation factor 1 α* gene), 100 U of T7 RNA polymerase (~0.83 μM), and the transcription buffer (40 mM Tris-HCl pH 8, 8 mM MgCl_2_, 2 mM spermidine-HCl, 25 mM NaCl). Vesicles were placed at 37 °C in a water bath under continuous illumination with a 860 nm/2.6 W LED lamp, and confocal microscopy images were recorded after 30, 60, 90 and 120 minutes of incubation. 5 μM Acridine Orange (AO) has been added in order to record the green fluorescence of the AO-RNA complex (λ_ex_ 488 nm, λ_em_ from 510 to 600 nm, peak at 530 nm).

Negative and positive control ASAPs samples were prepared similarly; the first by removing all nucleotides from the transcription kit, and the second by including ATP in the transcription kit. The AO response of the RNA-synthesizing transcription kit was verified in separated experiments carried out in bulk (negative control samples: buffer, or DNA template, or complete transcription kit lacking ATP; positive control sample: RNA from torula yeast (Sigma, #R6625)), and monitored by confocal micro-spectrofluorimetry (λ-scan).

### Confocal microscopy and quantitative image analysis

A confocal laser-scanning microscope Leica SP8 X was used for image acquisition. The emission signal from the fluorescent dyes used in this study was recorded by employing the usual settings (Nile Red: λ_ex_ 550 nm, λ_em_ 600-700 nm; pyranine: λ_ex_ 488 nm, λ_em_ 500-600 nm; AO-RNA complex: λ_ex_ 488 nm, λ_em_ 500-700 nm, peak at 530 nm). λ-Scans were performed with a HC PL APO CS2 20×/0.75 dry objective, recording fluorescence every 5 nm. Samples (30 μL) were placed in micro-wells plastic slides (ibidi GmbH, Gräfelfing-Münich, Germany, #81821). Image analysis was carried out by the free software ImageJ [M5].

