## Supplementary Information for "Light-driven ATP production promotes mRNA biosynthesis inside hybrid multi-compartment artificial protocells"

#### Table of Contents

Text S1. Terminology Details
Text S2. Details on chromatophore preparation
Text S3. Determination of RC amount and orientation in chromatophores
Text S4. Preliminary and control experiments about ATP-producing chromatophores
Text S5. Discussion on turnover number of ATP synthase
Text S6. Measuring ATP inside individual protocells
Text S7. Details of the minimal kinetic model

Table S1. Features of a 68 nm chromatophore
Table S2. Features of an average protocell (diameter: 20 μm)
Table S3. Numerical values used in the minimal kinetic model
Table S4. Composition of DMSZ medium (titrated with NaOH to final pH 6.9)
Table S5. Composition of the SL6 solution (pH 3-4)
Table S6. Composition of iron(II) citrate solution (pH 6.5), to be prepared under nitrogen

Figure S1. Cryo-EM images of chromatophores
Figure S2. Cryo-EM images of ATP synthases
Figure S3. ATP production assays of chromatophores in bulk solution
Figure S4. The droplet transfer method (details)
Figure S5. Increase of pH in the protocell free volume
Figure S6. Quantification of ATP produced inside protocells by entrapped chromatophores (OD 10)
Figure S7. Microspectrofluorimetry of acridine orange-mRNA complex

References of the Supplementary Information

**Text S1. Terminology Details**

In this manuscript, the term "protocells" is used with the meaning of "cellular prototypes", *i.e.* prototypes of compartments that mimic a cellular structure, behaviour or functionality. In this context, giant vesicles are, *per se*, "artificial protocells" since these compartments, prepared by a defined synthetic protocol, reproduce the size, the lipid composition and some properties of cell membranes, *e.g.* the metabolite passive transport across the lipid bilayer driven by a concentration gradient.
Artificial Simplified-Autotroph Protocells, or ASAPs, refer to protocells that mimic the capability of some living systems of sustaining their metabolism by exploiting an external energy source such as light and/or inorganic substrates: photoautotrophs and chemoautotrophs, respectively. The attribute "Simplified" emphasize that ASAPs are examples of artificial systems, inspired to living systems, attempting to reproduce the autotrophic capability at a proof-of-concept level.
Photoactive ASAPs that transduce light energy into chemical energy (ATP) and support DNA transcription are here presented.

**Text S2. Details on chromatophore preparation**

The production of membrane vesicles from bacteria can be achieved in different ways [S1, S2, S3]. In the case of *R. sphaeroides*, chromatophores result from cellular lysis, typically accomplished by mechanical treatments such as French press or sonication [S4, S5].

Sonication was ruled out because it can damage macromolecular and supra-macromolecular structures, and also lead to chromatophores with a low degree of orientation.

French press favours the formation of chromatophores due to the sudden pressure variation (from high to low), leading to cell explosion with a consequent rupture of cell wall, and a rearrangement of the cytoplasm membrane. The pre-existing intra-cytoplasmatic membrane invaginations quickly close in the desired "inside-out" orientation (everted vesicles), and efficiently entrap the solutes present in the periplasm (*i.e.* cytochrome $c_2$).

In this study, we opted for a single French press passage because multiple passages (which would increase the yield of chromatophores) would have been detrimental. In fact, chromatophores obtained from the first passage would break and reseal, losing their content and randomising the orientation of macromolecular machinery.

We expect, therefore, that a single French press passage will generate four types of structures:

*a.* "Functional" chromatophores. These are the large majority of chromatophores, in which RC and bc1 expose their cytochrome $c_2$ binding sites to the lumen, whereas the $F_1$ subunit of ATP synthase points outwards, to the external solution. The chromatophore lumen contains a sufficient amount of cytochrome $c_2$ (in reduced form, cyt$^{2+}$), and the membrane hosts the ubiquinone pool. In such an arrangement, chromatophores can be employed as ATP-producing photosynthetic organellae inside a larger vesicle. Importantly, they count as "functional" in the RC charge-recombination assay (Figure 2h-i).

*b.* Chromatophores with inside-out orientation of the membrane proteins, but lacking a sufficient amount of cyt$^{2+}$ in their lumen. These are actually functional chromatophore (*i.e.* are able to produce ATP in their external solution upon continuous illumination), but would count as "non-functional" in the RC charge-recombination assay, simply because RC in the charge-separated state cannot be reduced by cyt$^{2+}$, present at low concentration, and therefore unbound to RC in the moment of the light flash. Cryo-EM imaging cannot distinguish among chromatophores with high or low cytochrome $c_2$ concentration in their lumen.

*c.* Chromatophores with oppositely oriented membrane proteins (right-side out). These are "non-functional" structures in all conditions, and clearly count as "non-functional" in the RC charge-recombination assay. It should be noted, however, that their presence in the sample is not supported by cryo-EM imaging and therefore they are rarely generated by French press lysis. This observation agrees with the general knowledge on the production of membrane vesicles from bacteria [S1, S2, S3].

*d.* Open bilayer fragments carrying all – or parts – of the photophosphorylation machinery. These are "non-functional" structures in all conditions, and also count as "non-functional" in the RC charge-recombination assay. Their presence is confirmed by cryo-EM and cryo-electron tomography.

In conclusion, cryo-EM, cryo-electron tomography and the results from the RC charge recombination experiment (Figure 2) suggest that type *a* chromatophores ("functional" chromatophores) account for about 70% of all RC-containing structures. The remaining 30% accounts for *b*, *c* and *d* structures, where *c* is probably negligible, and we also expect that *b* >> *d*.

It should be noted that, strictly speaking, type *a* and *b* chromatophores are not necessarily distinct structures, because not all RCs in a chromatophore can interact with $cyt^{2+}$ in the exact moment of light flash. Ultimately, this parameter depends on the $cyt^{2+}$ concentration.

#### Text S3. Determination of RC amount and orientation in chromatophores

To determine the exact fraction of type *a* chromatophores (Text S2) out of the whole population of particles produced by French pressing, we probe RC orientation by charge recombination experiments.

Upon a saturating flash light excitation, a charge-separated state (CSS) is very rapidly produced in all RC molecules (~10 ps). In particular, the bacteriochlorophyll dimer (D),is oxidized and the photogenerated electron travels through RC, finally reaching a bound quinone (Q), which is reduced.

$$DQ + h\nu \rightarrow D^{+}Q^{-}\cdot$$

Note that D and Q are located in two distant sides of RC.

The transition from the ground state DQ to the CSS ($D^{+}Q^{-}\cdot$) brings about a reduction of absorbance at 860 nm (the absorption band of bacteriochlorophyll).

RC in the CSS can evolve in two ways:

- in the presence of enough $cyt^{2+}$ on the same side where $D^{+}$ is located, $cyt^{2+}$ is expected to be bound to RC and reduces $D^{+}$ ($cyt^{2+} + D^{+} \rightarrow cyt^{3+} + D$), so that RC quickly reaches a $DQ^{-}$ state ($\tau \sim 1\ \mu s$); such new state has the same absorbance of the ground state in the 860 nm region, and therefore no spectral changes are recorded in the time resolution of the instrument; RCs undergoing this path are silent in flash light excitation/charge recombination experiments;
- in absence of $cyt^{2+}$ on the same side where $D^{+}$ is located (or when $cyt^{2+}$ concentration is low), the CSS undergoes a charge recombination ($D^{+}Q^{-} \rightarrow DQ$). Charge recombination is slow ($\tau \sim 0.1\text{-}1$ s) and can be followed by the characteristic recovery of the ground state absorption band at 860 nm.

In conclusion, the initial amplitude of $\Delta A_{860}$, *i.e.* $\Delta A_{860}(0)$ (just after the light flash, before starting the charge recombination curve), is proportional to the amount of RC molecules in the CSS that cannot be reduced by $cyt^{2+}$. These will be RC molecules that are located in type *b*, *c*, or *d* structures (see Text S2).

When the sample is treated with detergent (1% w/v LDAO), chromatophores and bilayer fragments are transformed into micelles, and the entrapped $cyt^{2+}$ will be released in the external medium (~10,000-fold dilution). This means that no RC, after entering the CSS, can be reduced effectively by the very diluted $cyt^{2+}$, and thus *all* RC molecules will contribute to $\Delta A_{860}(0)_{,LDAO}$, undergoing a charge recombination pathway thereafter.

The total amount of RC in the sample can be obtained from $\Delta A_{860}(0)_{,\mathrm{LDAO}}$. Instead the amount of RC in type *a* chromatophores can be obtained from $[\Delta A_{860}(0)_{,\mathrm{LDAO}} - \Delta A_{860}(0)]$.

Chromatophores with OD (860 nm) = 0.6 were assayed (Figure 2h) of the main text). $\Delta A_{860}(0)$ = 1.8 mAU, corresponding to 17 nM RC; whereas $\Delta A_{860}(0)_{,\mathrm{LDAO}}$ = 5.5 mAU, corresponding to 52 nM RC. The latter value corresponds to 9 μM RC when OD = 100. About 70% of RC belongs to type *a* chromatophores. Therefore, the amount of RC embedded in type *a* chromatophores (OD 100) in a typical preparation is 6.3 μM.

Between-experiments reproducibility lies within the ±10% range.

**Text S4. Preliminary and control experiments about ATP-producing chromatophores**

A first series of measurements were carried out to optimize instrumental response with respect to the luciferin-luciferase luminescence assay, leading to the following "standard" conditions for the assays (referred to 10 μL reaction mix): $5.72 \times 10^{16}$ chromatophores/L (OD = 50), 2 mM ADP, 100 mM Pi (pH 8.0), and 3 minutes red light irradiation (860 nm). The screening data for assessing the ATP*syn* activity are shown in Figure S3a, where the following three factors were varied at binary yes/no levels: chromatophores, ADP + Pi, light. As expected, ATP synthesis occurs only when all factors are present (however, a small capacity of synthesizing ATP has been recorded when chromatophores, ADP and Pi were mixed in the dark, resulting in a 18% yield when compared to the complete set of factors).

Experiments at variable ADP concentration have shown that ATP is produced in good yield (60% conversion, Figure S3b) irrespective of ADP concentration (250-2000 μM). This is expected if ATP synthase is working at ~$V_{\max}$, a reasonable hypothesis, because in our conditions $[\mathrm{ADP}]_0 >> K_\mathrm{M}$ ($K_\mathrm{M}$ ~10 μM [S6]. The 60% conversion allows the calculation of the reaction quotient $Q_p$ for the phosphorylation reaction coupled with the proton translocation, *i.e.*

$$\begin{array}{lr} \mathrm{ADP} + \mathrm{P_i} \rightleftarrows \mathrm{ATP} + \mathrm{H_2O} & \Delta G'_p \\ n\mathrm{H}^+_{\mathrm{in}} \rightleftarrows n\mathrm{H}^+_{\mathrm{out}} & n\Delta\tilde{\mu}_{\mathrm{H}^+} \end{array}$$

An estimate of the proton motive force across the chromatophore membrane can be obtained according to the equations (1-3) shown below [S7]:

$$n\Delta\tilde{\mu}_{\mathrm{H}^+} = \Delta G^{o\prime}_p + RT\ln Q_p \quad (1)$$

$$Q_p = \frac{[\mathrm{ATP}]c^o}{[\mathrm{ADP}][\mathrm{P_i}]} \quad (2)$$

$$\Delta p = \frac{\Delta\tilde{\mu}_{\mathrm{H}^+}}{F} = \frac{\Delta G^{o\prime}_p + RT\ln Q_p}{nF} \quad (3)$$

where the symbols have their usual thermodynamic meaning, and $\Delta p$ is the proton motive force, expressed in V. Accordingly, when $\Delta G_p{}^{\circ\prime}$ = +30.5 kJ/mol [S8]; [ATP] = $1.2 \times 10^{-3}$ M; [ADP] = $0.8 \times 10^{-3}$ M; [Pi] = $98.8 \times 10^{-3}$ M; c° = 1 M, $n$ = 3, the calculated $Q_p$ is 15.2, and $\Delta p$ results to be 0.129 V at 25 °C.

We have verified the effect of sugars (glucose or sucrose) that need to be included in chromatophore-containing GVs because of the centrifugation step. Sugars were added to chromatophores before irradiation (200 mM glucose or 200 mM sucrose), or to chromatophores mixed with a known amount of ATP before performing the luciferin-luciferase assay. When compared with sugar-free assays, sugars reduce the final luminescence readouts of about −30%, possibly affecting both the photophosphorylation reaction and the luciferin-luciferase assay. These results prompted us to apply the standard addition method for the quantification of intra-GVs ATP concentration.

The effect of LDAO surfactant (from 0.001 to 1 % w/v in the 10 μL reaction volume) was also tested (to verify whether it can be used for releasing ATP from giant vesicles). LDAO, added after the ATP photosynthesis inside chromatophores, strongly reduces the luminescence readout (Figure S3c). These results prompted us to release the intra-GVs ATP by freezing-and-thawing rather than by detergent-induced GV lysis. Interestingly, similar inhibitory results have been reported in the case of other surfactants, like SDS or benzalkonium chloride [S9], whereas Triton-X-100 acts as an enhancer [S10].

#### Text S5. Discussion on turnover number of ATP synthase

Various values for the turnover number of ATP synthase are available in literature (but most of them refer to *E. coli* ATP synthase). Values in the 13-35 $s^{-1}$ range have been reported [S11, S12, S13, S14] but several studies have indicated higher rates under other conditions. Among the highest values, [S15] reported 50 $s^{-1}$ (23 °C), [S16] reported 90 $s^{-1}$ (24 °C), whereas very high turnover numbers have been reported when few active ATP synthase molecules were powered by large electrochemical proton gradients (e.g., 230 $s^{-1}$ and higher [S13]. Additional values are found in [S13], including those of mitochondria's and chloroplasts' ATP synthase. Single-molecule studies reveal that individual ATP molecules can display a rotation rate of 230 revolution/s at 23 °C (which corresponds to a turnover number of 690 $s^{-1}$ considering 3 synthesized ATP/revolution).

The turnover number of *Rhodospirillum rubrum* ATP synthase, according to early studies, is about 3.5 ATP/min $BChl^{-1}$ [S17]. The turnover number of ATP synthase in our *Rhodobacter sphaeroides* R26. chromatophores (OD 50) is of the same order of magnitude (1.7 ATP/min $BChl^{-1}$). On the other hand, more recent studies reported a $k_{cat}$ of 28 $s^{-1}$ for *Rhodobacter sphaeroides* chromatophores [S6], whereas in this study we measured a $k_{cat}$ of 100 $s^{-1}$.

#### Text S6. Measuring ATP inside individual protocells

The luciferin-luciferase assay does not allow the determination of ATP production inside individual protocells. At this aim, microscopy imaging could be used if ATP-responsive fluorescent probes are employed. *Ad hoc* designed rhodamine derivatives have been reported as ATP sensors [S18, S19]. Unfortunately, the responsiveness of these probes was not satisfactory in our conditions, even in bulk experiments.

It should be noted that Pols *et al.* [S20] have recently reported the use of Perceval HR [S21], an ADP/ATP ratio protein sensor, to monitor the ADP to ATP conversion inside conventional sub-micrometre vesicles.

#### Text S7. Details of the minimal kinetic model

The ordinary differential equation set, that describes the time evolution of the mRNA transcription, sustained by driven-light ATP formation, is as follows:

$$\frac{d}{dt}F_{ADP} = -k_{ATPsyn}^{ps}\frac{F_{ADP}}{K_{M_{ADP}}^{ps} + F_{ADP}}$$

$$\frac{d}{dt}F_{ATP} = +k_{ATPsyn}^{ps}\frac{F_{ADP}}{K_{M_{ADP}}^{ps} + F_{ADP}} - k_{RNApol}^{ps}\frac{F_{ATP}}{K_{M_{ATP}}^{ps} + F_{ATP}}$$

$$\frac{d}{dt}F_{RNA} = +k_{RNApol}^{ps}\frac{F_{ATP}}{K_{M_{ATP}}^{ps} + F_{ATP}} - k_{dec}^{ps}F_{RNA}$$

In particular, this simple ODE set describes the process as two consecutive steps: 1) the transformation of ADP in ATP and 2) its polymerization. The first step, the conversion of the ADP in ATP is driven by the light energy while the polymerization of nucleotides is sustained by the ATP freshly produced. ADP is a reagent in defect and, therefore, ATP is the species that mainly determines the mRNA polymerization rate. Indeed, all the other bases are considered in strong excess so that their concentration is greater than the relative $K_M$ and they do not influence the rate. The calculated time course of the mRNA fluorescence has been then optimized in order to get both the pseudo kinetic constants, $k^{ps}_{ATPsyn}, k^{ps}_{RNApol}$ and $k^{ps}_{dec}$, and Michaelis-Menten constants, $K^{ps}_{M_{ADP}}$ and $K^{ps}_{M_{ATP}}$, respectively, all involved in the kinetic model.

In particular, the pseudo kinetic constants contain the concentration of all the species that participate to each kinetic step, even if they remain constant throughout the process. DNA, T7 RNA polymerase and ATP synthase can be considered constant along with species in large excess but at a concentration value comparable to the relative $K_M$, like phosphate.

In order to convert fluorescence intensities in concentrations of the different species and then to make the best fit constants comparable with those reported in literature, some assumptions are necessary:

1. all the ADP molecules present in the system are converted in ATP and then polymerized with a yield of 100%.
2. the fluorescence intensity obtained by the formation of the complex between the Acridine-Orange and mRNA is proportional to the amount of genetic polymer formed and then directly related to the amount of polymerized ATP.

These two assumptions make possible to convert the highest value of the fluoresce obtained after 90 minutes (20.8 a.u.) to the initial concentration of the ADP $2.0 \times 10^{-4}$ M, while in order to convert this concentration into the maximum concentration of the mRNA chains we must divided this value for the average number of ATP contained in each mRNA strand: 576 bases.

In the Figure 4c, it is reported the comparison between the experimental data (point with error bars) and the best fit curves (solid lines) obtained by converting the intensity of fluorescence in concentration, according to assumptions 1 and 2.

To estimate the real kinetic constants from the best fit values, it is necessary to explicit the pseudo constant value formula:

$$k^{ps}_{ATPsyn} = \varphi k_{ATPsyn}[ATPsyn]\frac{[Pi]}{K_{M_{Pi}} + [Pi]}$$

$$k^{ps}_{Pol} = \varphi k_{RNApol}[Polym]\frac{[DNA]}{K_{M_{DNA}} + [DNA]}$$

$$k^{ps}_{dec} = \varphi k_{dec}$$

Where $[ATPsyn] = 1.3 \times 10^{-8}$ M, $[Pi] = 0.01$ M, $[Polym] = 8.32 \times 10^{-7}$ M and $[\mathrm{DNA}] = 12.53 \times 10^{-9}$ M are the concentrations of ATP synthase, phosphate, T7 RNA polymerase and DNA respectively, while $K_{M\ Pi}$ and $K_{M\ DNA}$ are the relative Michaelis-Menten constants for phosphate and DNA. $\varphi = 20.8/0.0002$ $M^{-1}$ is the conversion factor from fluorescence intensity to concentration for ADP and ATP.

The formula to convert the Michaelis-Menten constants are instead simpler:

$$K^{ps}_{M_{ADP}} = \varphi K_{M_{ADP}}$$

$$K^{ps}_{M_{ATP}} = \varphi K_{M_{ATP}}$$

#### Tables

**Table S1. Features of a 68 nm chromatophore**

| *Property* | *Value* | *Note* |
|---|---|---|
| Diameter (DLS) | 68 nm (mode) | From the number-weighted size distribution; the mean is $80 \pm 23$ nm. |
| Internal volume | $1.13 \times 10^5$ nm$^3$ | |
| Zeta-potential | $-30.9 \pm 0.6$ mV | Measured in this study. |
| Average fraction of anionic lipids | 60 mol% | From [S22, S23]. |
| RC | 33 | Calculated from the known value of RC density for chromatophores from *R. sphaeroides* grown in high-light conditions (ca. 2,300 RC/μm$^2$) [S24, S25]. |
| LH1 | 33 | From RC:LH1 1:1 [S26, S27]. |
| Number density of chromatophores (OD 100) with the photophosphorylation machinery in an inside-out orientation | $1.14 \times 10^{17}$/L | Calculated from the RC concentration (6.3 μM), determined by flash light/charge recombination experiments, and the known RC density for chromatophores derived from *R. sphaeroides* grown under high light conditions (ca. 2,300 RC/μm$^2$) [S24, S25].<br><br>According to our measurement, it also corresponds to [BChl] = 468 μM |
| Chromatophores captured volume (OD 100) | 1.3% | (*i.e.* 13 μL/mL) |
| bc1 | 5.5 | Calculated from the known RC/bc1 molar ratio 6/1 [S26]. |
| ATP synthase | 0.73<br><br>[1.6] | Estimated from the measured ATP synthase density *via* cryo-EM* and cryo- tomography, resulting 50 ATP*syn*/μm$^2$.<br><br>The value in square bracket is the measured number of ATP*syn*/chromatophore (irrespective of their size). |

(*) The value is best considered an underestimate because it does not keep into account the non-equatorial ATP*syn* that are possibly present in the chromatophores sampled for the measurement. Typically, most studies refer to 1 ATP*syn* /chromatophore (88 ATP*syn* /μm$^2$ if a 60 nm chromatophore is considered).

**Table S2. Features of an average protocell (diameter: 20 μm)**

| *Property* | *Value* | *Note* |
|---|---|---|
| Lipid | POPC | |
| Lipid concentration | 150 μM | Determined from size distribution and entrapment efficiency (with the hypotheses of spherical shape and unilamellarity). The value matches unpublished results from our laboratory, based on lipid quantitation by the Stewart assay. |
| Diameter | 20 μm | Determined by image analysis of 950 protocell |
| Efficiency of droplet transfer | 35% | Determined by measuring the concentration of chromatophores released in the O-solution due to failed w/o droplet-to-giant vesicle transfer |
| Number of protocell/mL | $2.2 \times 10^7$ | When the protocell pellet is suspended in 50 μL |
| Protocell captured volume | 14% | Total protocell volume/volume of external solution (*i.e.* 140 μL/mL) |
| Number of chromatophores (OD 10) entrapped in one 20 μm protocell | 48,000 | |
| Protocell volume occupied by chromatophores | 0.2% | |
| Protocell free inner volume | 99.8% | |

**Table S3. Numerical values used in the minimal kinetic model**

| | | *Value* | | *"set" or "fitting"* | *Comment* |
|---|---|---|---|---|---|
| *Reaction 1. Photophosphorylation* | | | | | |
| ADP | $[ADP]_0$ | 200 | μM | set | Experimental value |
| Pi | $[Pi]_0$ | 10 | mM | set | Experimental value |
| ATP*syn* | $C_{0,E1}$ | 13.8 | nM | set | Calculated from Table S1, if OD = 10 |
| ATP*syn* | $K_M$ (ADP) | 128 | μM | fitting | *Cf.* 10 μM [S13, S6];100 μM [S14] |
| ATP*syn* | $K_M$ (Pi) | 4.2 | mM | set | [S14] |
| ATP*syn* | $k_{cat}$ | 12.9 | $s^{-1}$ | fitting | 13-35 $s^{-1}$ [S11, S12, S13, S14] |
| *Reaction 2. Transcription* | | | | | |
| GTP, CTP, UTP | $[NTP]_0$ | 7.5 | mM | set | From the TX kit |
| DNA template | $C_{0,DNA}$ | 12.5 | nM | set | From the TX kit |
| T7 RNA polymerase | $C_{0,E2}$ | 0.832 | μM | set | From the TX kit |
| T7 RNA polymerase | $K_M$ (DNA) | 5 | nM | set | Taken from [S28, S29] |
| T7 RNA polymerase | $K_M$ (NTP) | 60.5 | μM | fitting | *Cf.* 80 μM [S28, S29] |
| T7 RNA polymerase | $k_{cat}$ | 0.13 | $s^{-1}$ | fitting | *Cf.* 0.482 $s^{-1}$ [S30]; 1.67 $s^{-1}$ [S28, S29]; 1 $s^{-1}$ [S31]; 2.2 $s^{-1}$ [S32] |
| *Reaction 3. RNA decay* | | | | | |
| | $k_{decay,mRNA}$ | negligible | | fitting | $7.9 \times 10^{-5}$ $s^{-1}$ (referred to polymerised nucleotide) [S28, S29]. |

**Table S4. Composition of DMSZ medium (titrated with NaOH to final pH 6.9)**

| Compound | Concentration | Units |
|---|---|---|
| $KH_2PO_4$ | 0.5 | g/L |
| $MgSO_4 \cdot 7\ H_2O$ | 0.8 | g/L |
| NaCl | 0.4 | g/L |
| $NH_4Cl$ | 0.4 | g/L |
| $CaCl_2 \cdot 2\ H_2O$ | 0.05 | g/L |
| D,L-malic acid | 1.5 | g/L |
| Yeast exctract | 2 | g/L |
| *p*-aminobenzoic acid | 0.001 | g/L |
| SL6 solution | 1 | mL/L |
| iron citrate solution | 6 | mL/L |
| ethanol | 1 | mL/L |

**Table S5. Composition of SL6 solution (pH 3-4)**

| Compound | Concentration | Units |
|---|---|---|
| $ZnSO_4 \cdot 7\ H_2O$ | 0.1 | g/L |
| $MnCl_2 \cdot 4\ H_2O$ | 0.03 | g/L |
| $H_3BO_3$ | 0.3 | g/L |
| $CoCl_2 \cdot 6\ H_2O$ | 0.2 | g/L |
| $CuCl_2 \cdot 2\ H_2O$ | 0.01 | g/L |
| $NiCl_2 \cdot 6\ H_2O$ | 0.02 | g/L |
| $Na_2MoO_4 \cdot 2\ H_2O$ | 0.03 | g/L |

**Table S6. Composition of iron(II) citrate solution (pH 6.5), to be prepared under nitrogen.**

| Compound | Concentration | Units |
|---|---|---|
| $FeSO_4$ | 0.608 | g/L |
| $Na_2SO_4$ | 0.528 | g/L |
| citric acid | 1.680 | g/L |

### Figures

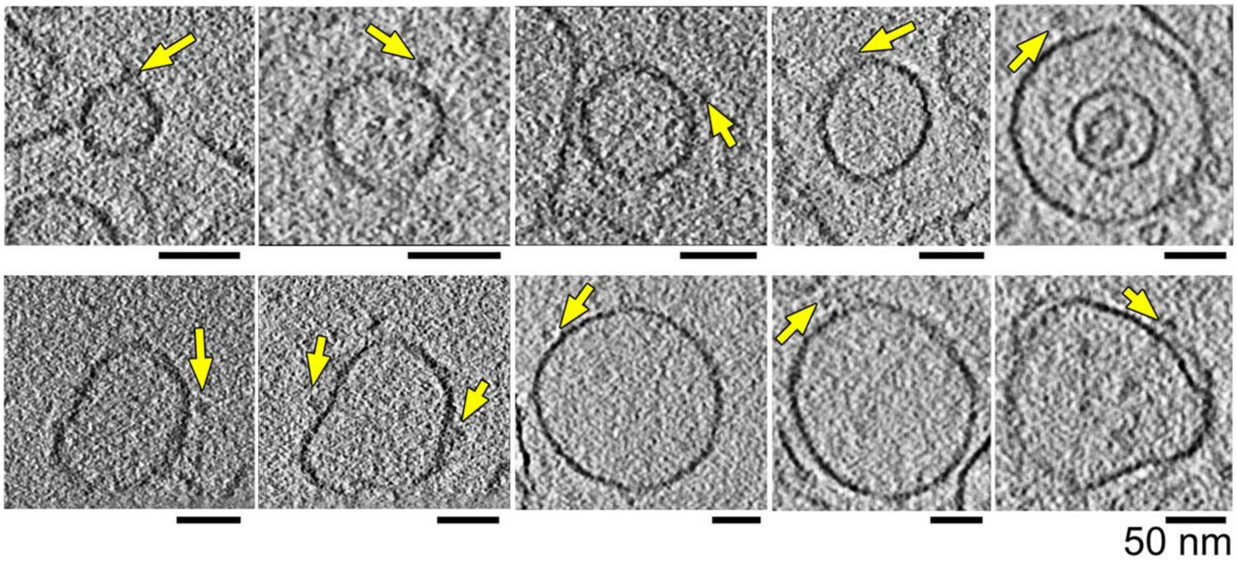


**Figure S1. Cryo-EM images of chromatophores.** Averaged cryo-tomographic slices of chromatophores vesicles carrying ATP synthase complexes (arrowhead). Size bars represent 50 nm.

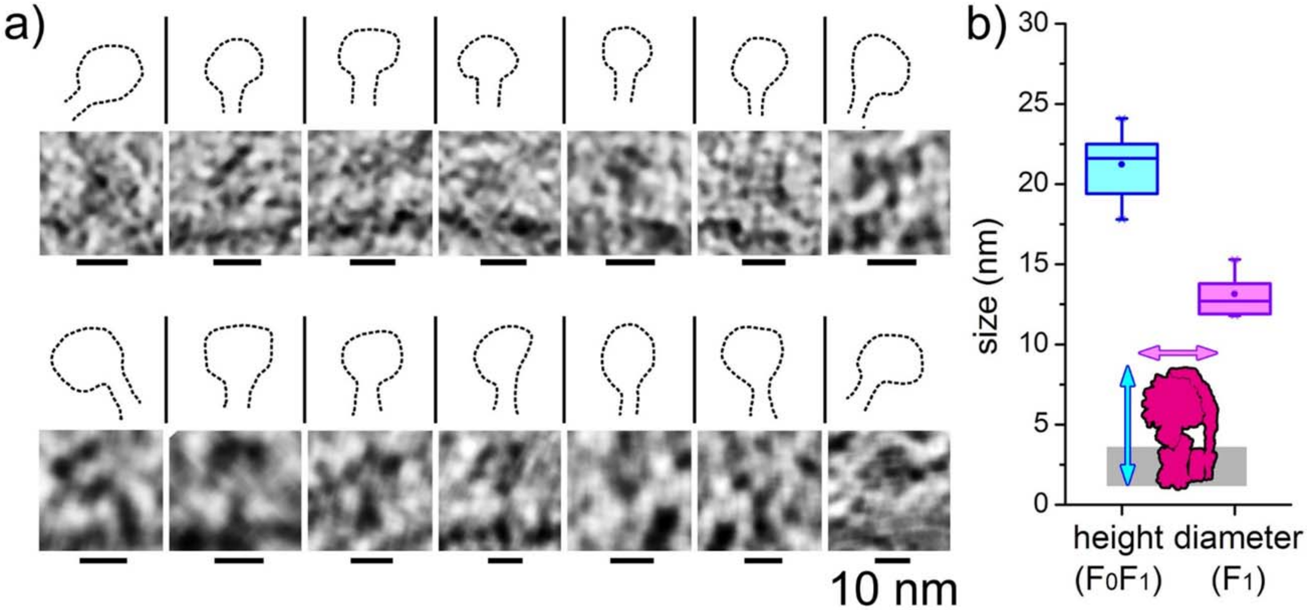


**Figure S2. Cryo-EM images of ATP synthases.** (a) Averaged central cryo-tomographic slices showing ATP synthase complexes present on the membrane of the analysed chromatophores. Scale bars represent 10 nm. (b) Average height (mean ± SD, 21.2 ± 1.9 nm) and diameters (mean ± SD, 13.2 ± 1.2 nm) as obtained by direct measurements ($n$ = 14). The horizontal lines of the boxes represent the first, the second, and the third quartiles. Reference values (referring to *E. coli* ATP*syn*) are 20 and 10 nm [S33, S34].

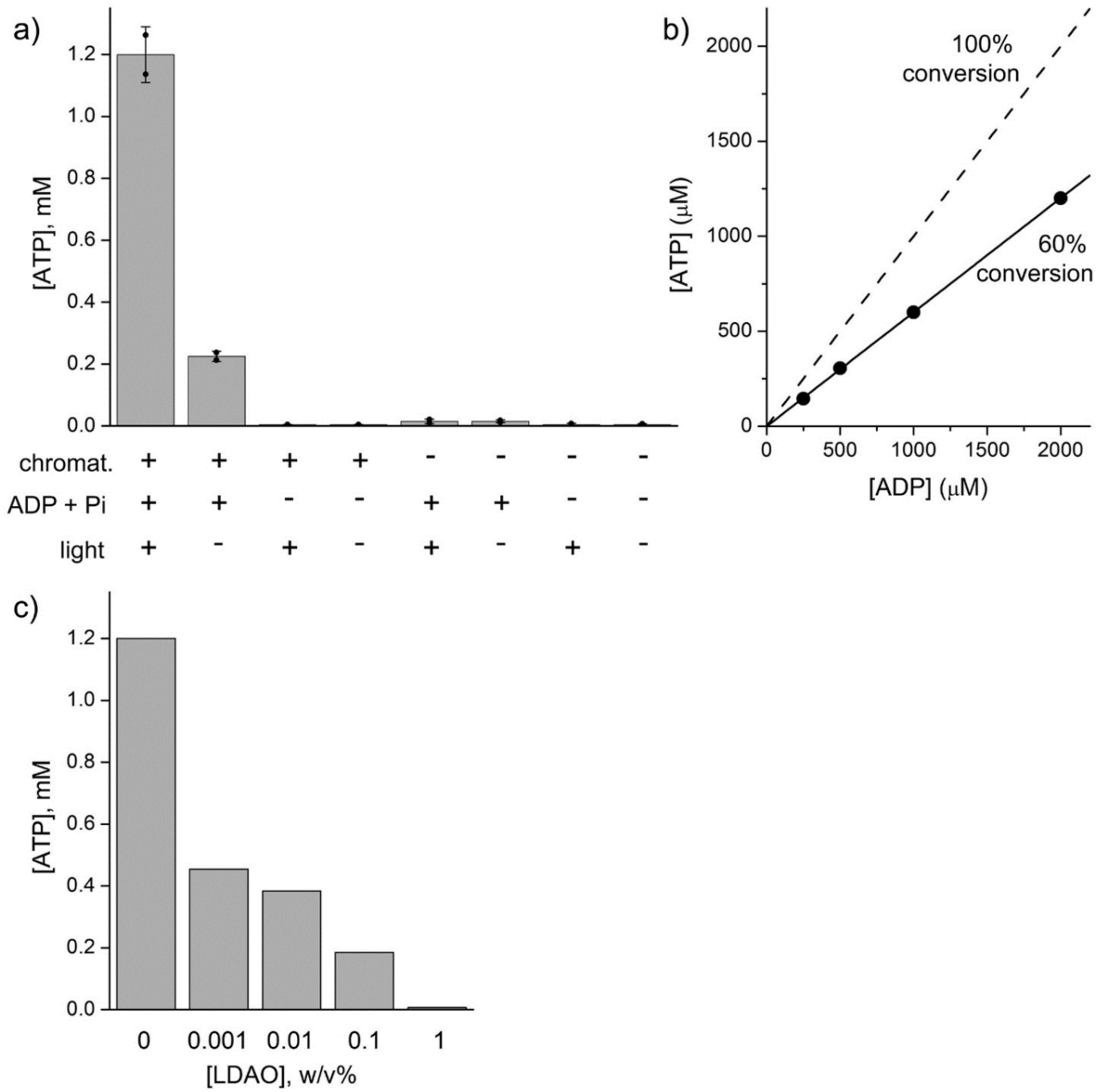


**Figure S3. ATP production assays of chromatophores in bulk solution.** (**a**) Factor testing. Eight samples containing chromatophores (OD 50), ADP (2 mM) and Pi (10 mM) were prepared according to the table below panel (a) and illuminated with 860 nm light for 3 minutes. Data are reported as mean ± SD of $n$ = 2 independently prepared samples. (**b**) Experimental measure of ATP produced by chromatophores by varying the externally added ADP concentration (250-2000 μM, referred to 10 μL of reaction mixture). Note the 60% conversion is essentially independent from ADP concentration. Being Pi in excess, this suggests that the position of the equilibrium is primary controlled by the proton motive force ($\Delta p$) across the chromatophore membrane. (**c**) Effect of LDAO on the assay. All concentrations (including ATP and LDAO) refer to10 μL reaction mixture (before dilution to 100 μL with 90 μL assay solution).

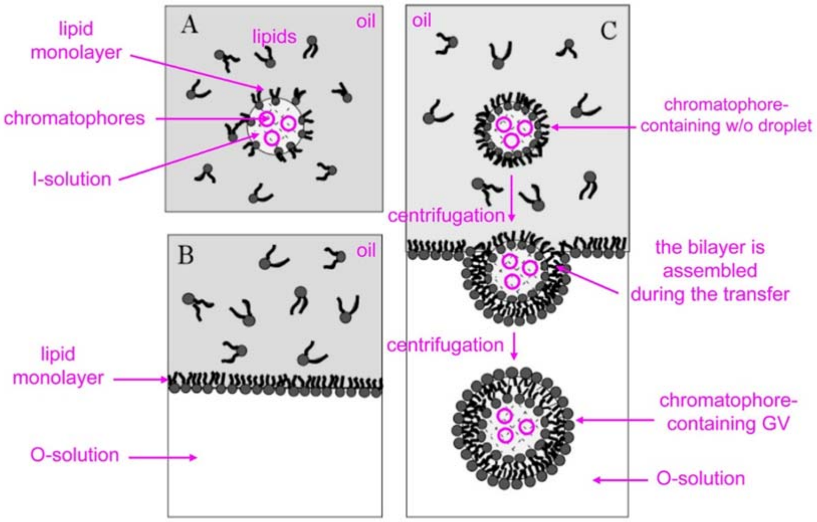


**Figure S4: The droplet transfer method (details).** Reproduced and adapted from [S35] ©2003 American Chemical Society (in particular, the original figure lacks all elements here coloured in magenta), with permission. The drawing is not to scale.

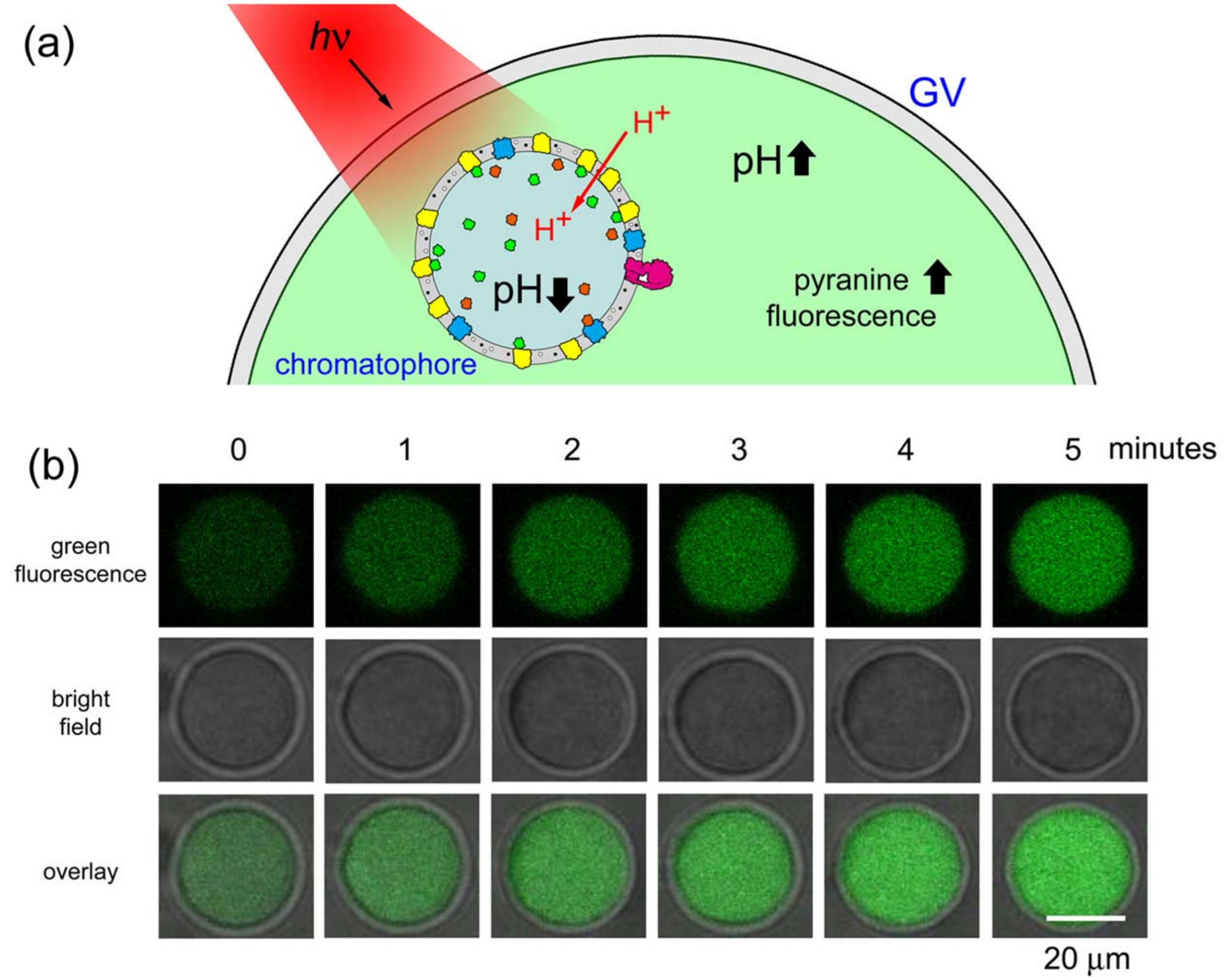


**Figure S5. Increase of pH in the protocell free volume as a result of light-induced proton translocation into the chromatophore lumen**. (**a**) *R. sphaeroides* chromatophores are encapsulated inside ~20 µm protocells made of POPC (1-palmitoyl-2-oleoyl-*sn*-glycero-3-phosphocholine) and illuminated with 860 nm light. Pyranine (trisodium 8-hydroxypyrene-1,3,6-trisulfonate) a membrane non-permeable pH-sensor, whose fluorescence increases along with pH increase, is co-encapsulated. In absence of ADP + Pi, illuminated chromatophores just translocate protons (from protocell lumen to the chromatophore lumen), so that the intra-chromatophore pH decrease, whereas the pH of the inner protocell volume increases, as revealed by the pyranine green fluorescence. (**b**) Confocal microscopy images. Fluorescence increases at a rate of 6.0 ± 1.8 a.u./min (mean ± SD, $n = 4$), which is statistically significant (two-tail $p < 0.05$) when compared to the negative control sample (1.8 ± 0.1 a.u./min, mean ± SD, $n = 12$). Protocells have been constructed by co-encapsulating chromatophores (OD 50) and pyranine (5 µM), in absence of ADP and Pi. Upon continuous irradiation, the RC-bc1 machinery (by means of $Q/QH_2$ and $cyt^{2+}/cyt^{3+}$ carriers), translocates protons from the protocell lumen into the chromatophore lumen, resulting in a pH increase of the intra-protocell solution. Negative control experiment (pyranine-containing protocells).

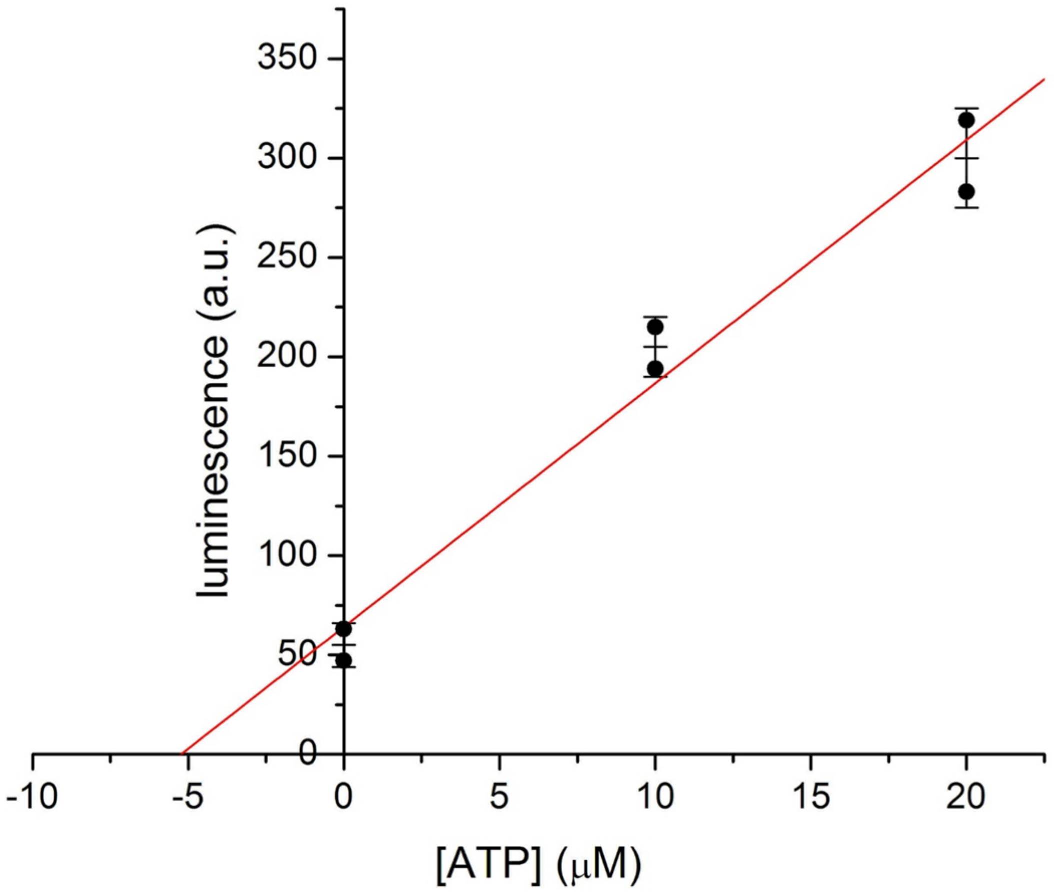


**Figure S6. Quantification of ATP produced inside protocells by entrapped chromatophores (OD 10).** The standard addition method was applied, by adding known amount of ATP to the samples. ATP concentrations refer to 10 μL reaction mixture (before dilution to 100 μL with 90 μL assay solution). Data are reported as mean ± SD of $n = 2$ independently prepared samples. The best-fit straight line is $y = (64 \pm 20) + (12 \pm 2)\ x$, intercepting the $x$-axis at 5.3 ± 1.8 μM. From the latter value, an average intra-protocell ATP concentration of ~38 μM can be estimated resulting from the concentration of released ATP (5.3 μM) and the protocell captured volume (14%, see Table S2).

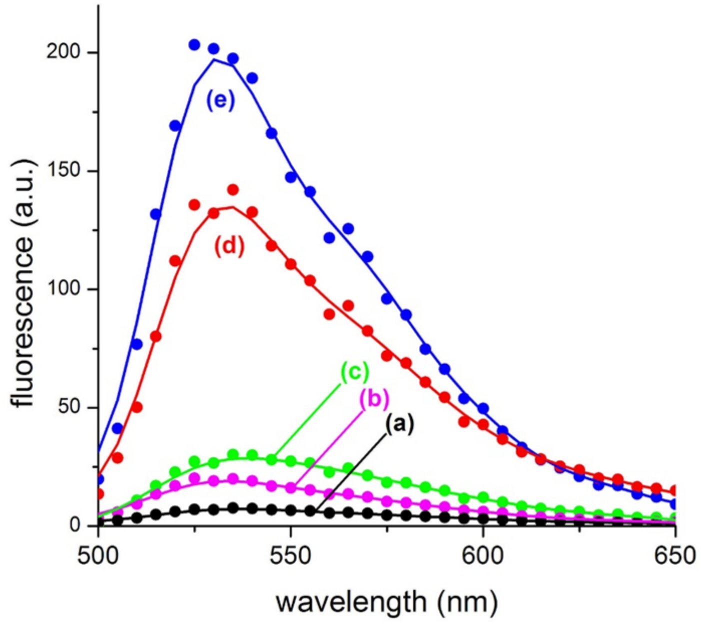


**Figure S7. Microspectrofluorimetry of acridine orange-mRNA complex.** The figure shows five fluorescence emission profiles as recorded by confocal microscopy by the λ-scan mode (excitation: 488 nm). In all experiment, carried out in bulk solution, 5 µM acridine orange (AO) was used. (*a*) AO + transcription buffer; (*b*) AO + transcription kit (-ATP); (*c*) AO + DNA template (0.5 µg in 20 µL); (*d*) AO + full transcription kit (including ATP) + DNA template (0.5 µg in 20 µL), incubated for 1 hour at 37 °C; (e) AO + 0.5 g/L torula yeast RNA extract.
